# Electrophysiological profiling of hiPSC-derived neurospheres using a novel NeuroMPS with integrated electrodes

**DOI:** 10.64898/2026.06.30.735615

**Authors:** Fulya Ersoy, Paolo Cesare, Lisa-Marie Erlandsdotter, Matthijs van der Moolen, Andrea Lovera, Sascha Mommo, Peter D. Jones, Peter Loskill

## Abstract

Despite advances in microphysiological systems, in vitro platforms for neuronal models remain limited by insufficient electrophysiological resolution, poor structural compatibility with 3D tissue architecture, and an inability to capture functional network dynamics alongside morphological and metabolic readouts. Here, we develop a neuromicrophysiological system (NeuroMPS) that pairs human iPSC-derived neurospheres, comprising neurons and glial cells, with tailored microelectrode arrays (MEAs) for non-invasive, high-resolution monitoring of neuronal network dynamics and functional maturation in vitro. The NeuroMPS combines a custom MEA with capped electrodes optimized for neurite-level signal detection and a glass microwell module that provides structural confinement and optical compatibility for imaging. Importantly, this configuration enables stable, longitudinal electrophysiological recordings from 3D neural constructs while supporting multimodal analyses. Following exposure to pharmacological modulators (PTX, TTX, bicuculline, CNQX, and 4-AP) and the neurotoxin rotenone, NeuroMPS detects alterations in network activity within minutes, even at the lowest concentrations tested, whereas morphological and metabolic changes emerge only at higher doses and later time points. This work provides a physiologically relevant, scalable, non-invasive platform that integrates high-sensitivity electrophysiological readouts with morphological and metabolic profiling to enable early prediction of compound-induced effects in human iPSC-derived 3D neural networks, with applications in neuropharmacology, neurotoxicology, and disease modeling.

## Introduction

In our aging society, neurodegenerative disorders such as Alzheimer’s disease and Parkinson’s disease represent a major global health burden and constitute the second leading cause of death in Europe[1]. Their multifactorial etiology and the inherent complexity of brain structure and function pose substantial challenges for developing effective therapeutics. Conventional experimental platforms, ranging from 2D in vitro models to animal models, often fail to recapitulate crucial human-specific neurophysiological dynamics, resulting in limited translational power and poor predictive accuracy for patient responses[2, 3].

Technological advances over recent decades have driven the emergence of 3D microphysiological systems (MPS) that more faithfully recapitulate human brain architecture and function in vitro[4–6]. Merging approaches from molecular and cellular neuroscience, bioengineering, materials science, and physics, these systems recreate key aspects of neural microenvironments, including multicellular composition, 3D cytoarchitecture, and dynamic cell–cell interactions[7, 8]. As a result, 3D MPS platforms surpass the biological relevance and functional complexity achievable with traditional monolayer cultures.

3D neuronal systems, despite their inherent constraints, offer substantial potential for recapitulating cerebral processes with heightened fidelity[9, 10]. The integration of an extracellular matrix, although technically demanding, represents a surmountable barrier. Hydrogel scaffolds such as Matrigel provide ECM-like structural support and have been shown to improve the reconstruction of physiologically relevant neural tissue environments[11, 12]. As an alternative to hydrogel - embedded cultures, multicellular assemblies, including neurospheres and brain organoids, provide self-organized 3D architectures derived from iPSCs[13–15] or NPCs[16], subsequently differentiating into diverse neuronal and glial lineages. These constructs exhibit a degree of cellular and architectural complexity approaching in vivo conditions, rendering them particularly advantageous for interrogating neuronal network dynamics. Neurospheres enable the controlled examination of disease-associated phenotypes while preserving essential microcircuit features[17, 18] whereas more elaborate organoid systems, such as cerebral and midbrain organoids, facilitate the exploration of central nervous system organization and functional maturation[19, 20].

Although all 3D neuronal platforms contribute meaningfully to the study of neuropathophysiological mechanisms, assembled cellular constructs (neurospheres and organoids) occupy a unique position at the interface between in vitro and in vivo neurobiology, characterized by advanced cellular composition, developmental progression, and tissue-like architecture. Nonetheless, extracting quantitative readouts from these systems remains more constrained than from hydrogel-supported cultures. Morphological assessment of organoids and neurospheres is technically challenging, and obtaining functional information from these dense structures is similarly limited. Recent advances in imaging, optical clearing, and volumetric analysis are progressively improving data quality. Furthermore, 3D cultures frequently experience insufficient or heterogeneous nutrient distribution, resulting in elevated core cell death[21]. This limitation is increasingly mitigated through refined engineering strategies[22, 23], which enhance nutrient diffusion and overall tissue viability. Despite these advances, direct readout of electrical activity, the most informative measure of neuronal function, remains a major bottleneck. Although several sophisticated devices, such as shank-type[5] and shell-type microelectrode arrays (MEAs) [24] that probe volumetric network activity, have recently been introduced to capture electrophysiological signals from 3D cultures. However, these platforms are often invasive to culture, since electrodes are inserted into the 3D architecture, are high-maintenance, lack throughput, and are incompatible with the requirements of automated culture workflows. These limitations have hindered the broader adoption of 3D electrophysiological assays in drug discovery and disease modeling.

We present a NeuroMPS integrating custom glass-based MEA within a microplate-compatible format to enable scalable, high-fidelity electrophysiological interrogation of 3D neuronal circuits. The platform combines iPSC-derived neurospheres with microfluidic compartmentalization to support multimodal characterization, including high-resolution imaging, molecular readouts, and non-invasive electrophysiology. Neurospheres cultured in this system were evaluated across key functional domains, viability, neuronal maturation, electrophysiological activity, and pharmacological responsiveness, alongside a neurotoxin assay demonstrating the robustness of electrical readouts. Together, these results establish a unified and physiologically representative MPS framework suited for automated, higher-throughput workflows, advancing both fundamental neurobiology and translational research.

## Results

### Multimodal NeuroMPS enables integration of 3D neuronal models

NeuroMPS is a user-friendly, multimodal platform for studying 3D neurospheres in vitro at functional, morphological, and metabolic levels. The system combines a specialized MEA featuring capped microelectrodes (CMEs) with a monolithic quartz-glass module containing nine individual wells, providing electrophysiological capabilities and optical accessibility (Figure 1a). Neurospheres can be placed in the wells suspended on dedicated pins positioned 300 µm above the well bottom. The spheres are surrounded by 12 µl of gel matrix and 180 µl of medium (Figure 1c). Its open-well design allows easy supernatant collection, supporting downstream biochemical analyses. The specialized CMEs enable non-invasive electrophysiological recordings from 3D cultures. The MEA design follows the footprint of commercially available 120-electrode MEAs (MEA2100-System, Multi Channel Systems GmbH, Reutlingen, DE), with each of the 9 wells containing 13 recording electrodes and 1 reference electrode (Figure 1a-b). The CMEs consist of an electrode, microtunnels, and a sealing cap, fabricated using photolithography techniques[7] (Figure S1). Each well features a tissue compartment with small glass pins located 300 µm above the MEA bottom. These pins ensure accurate placement of neurospheres, while allowing neurite extension through the gel matrix and into the CMEs (Figure 1b). The neurites arborize into the gel matrix and grow in all directions in the tissue chamber (Figure 1d). The culture media above the tissue compartment provide oxygen and nutrients to the dense neuronal network. The microwell module is designed with the footprint of a standard 96-well plate, facilitating integration into automated workflows (Figure 1a). This architecture supports non-invasive, long-term monitoring of neuronal tissue, enabling longitudinal functional, morphological, and metabolic assessments.

**Figure 1.**
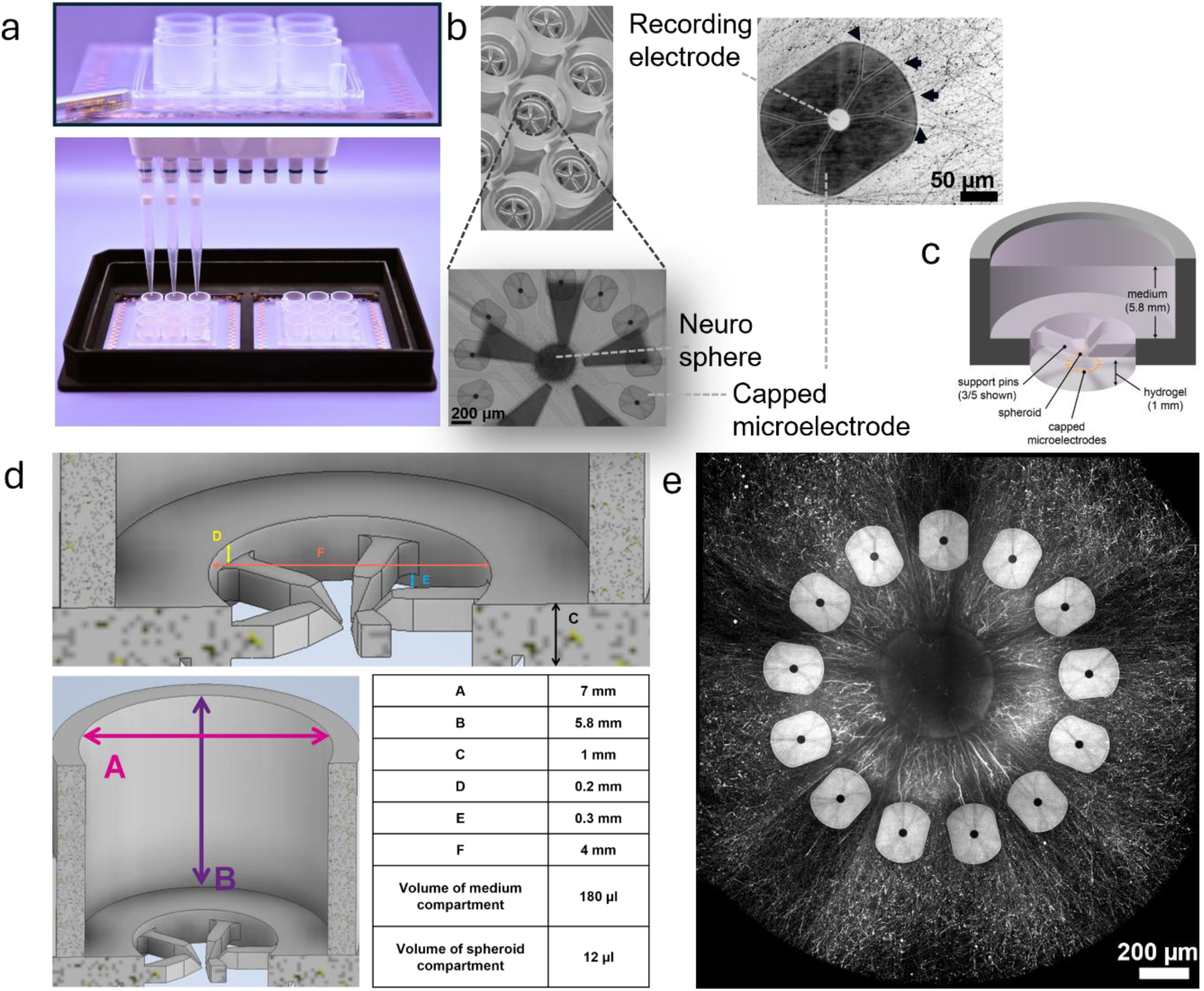
NeuroMPS with integrated electrodes for neurosphere culture and recording. **a** The NeuroMPS comprises nine wells and adheres to ANSI/SLAS microplate standards, ensuring compatibility with multi-pipette workflows and automated handling systems. The custom module follows a 96-well plate footprint and accommodates two devices. **b** Each well contains five pins for stable neurosphere positioning and incorporates 13 CMEs with microtunnels that guide neurites toward the recording electrode (black arrows). **c** Schematic of a cross-sectional view of one well. **d** Schematic illustrating the dimensional layout of the tissue and media compartments within each well. **e** Representative confocal image of a GFP-transduced neurosphere showing neurite outgrowth toward the CME structures.

### hiPSC-derived neurospheres incorporate functional neuronal and glial populations

The NeuroMPS was developed to facilitate the study of assembled 3D neuronal cultures. Here, we employed neurospheres generated from hiPSC-derived neuroprogenitor cells (NPCs) using a modified protocol adapted from Pamies et al.[16], which comprises spheroid formation, differentiation, and maturation over a period of eight weeks under continuous orbital shaking (Figure 2a).

**Figure 2.**
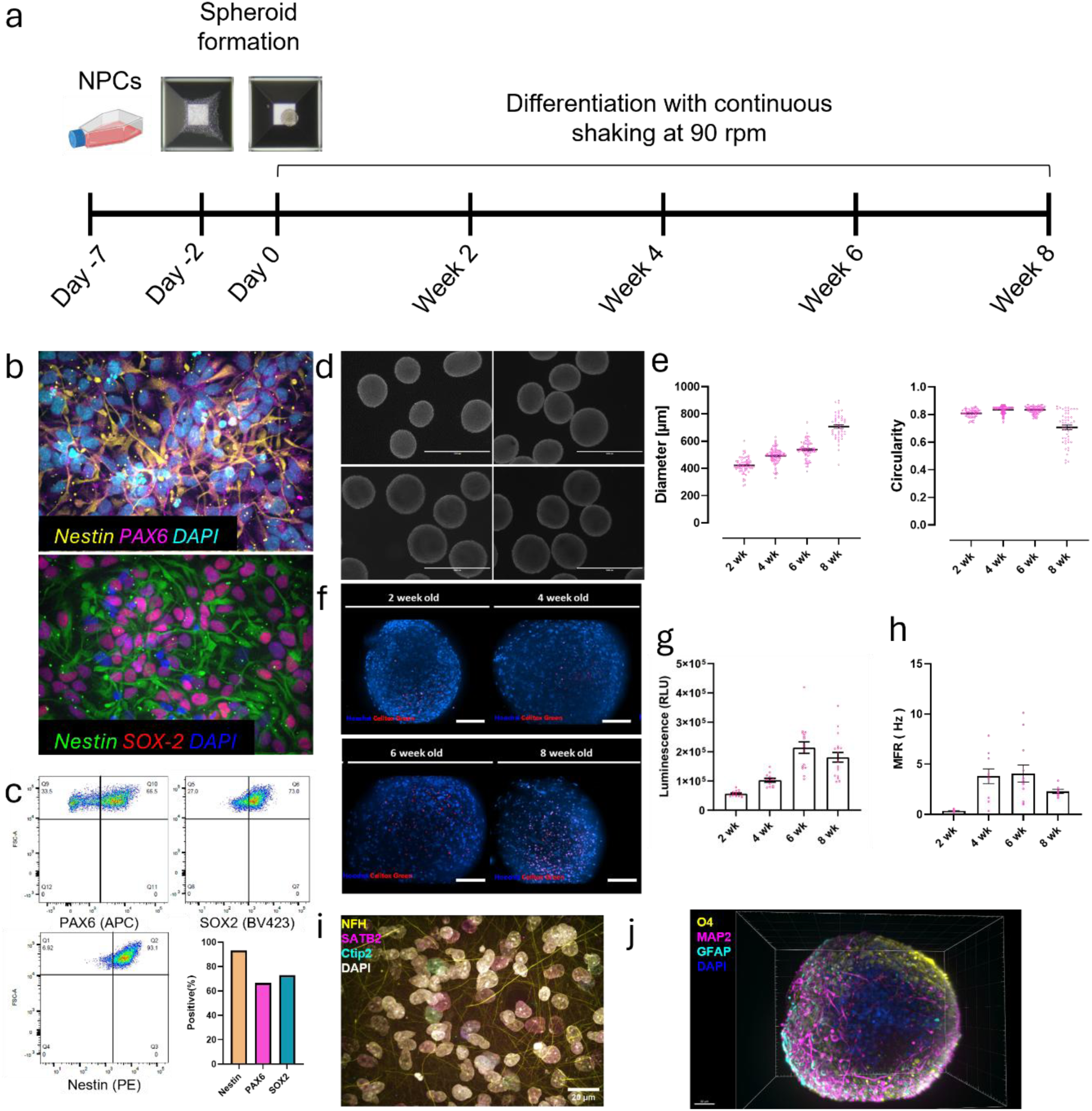
Characterization of neurospheres derived from NPCs. **a** Schematic timeline of neurosphere generation from NPCs. **b** Immunocytochemistry of KOLF2.1J NPCs showing co-localization of nestin with PAX6 (top) and SOX2 (bottom); nuclei were counterstained with DAPI. Scale bars, 50 µm. **c** Flow cytometry analysis of KOLF2.1J NPCs indicating that >60% of cells are positive for PAX6, SOX2, and nestin. **d** Representative brightfield images of neurospheres. Scale bar, 1 mm. **e** Quantification of neurosphere diameter and circularity over time (N > 50); lines indicate mean values. Statistical analysis was performed using one-way ANOVA; asterisks denote significant differences over time. **f** Representative live imaging of neurospheres at weeks 2, 4, 6, and 8; nuclei are labeled with Hoechst (blue) and dead cells with CellTox™ Green (red). Scale bar, 100 µm. **g** Viability assessed by ATP-dependent luminescence (N = 12–18), showing an increase up to week 6 followed by a decline at week 8. **h** Development of electrophysiological activity over 8 weeks of differentiation (8–10 neurospheres per time point; bars represent mean values). **i** Immunostaining of 6-week-old neurospheres for neurites (NFH), cortical layer V–VI marker (CTIP2), and upper-layer marker (SATB2), with DAPI nuclear counterstain; scale bar, 50 µm. **j** Neurospheres express mature neuronal markers for dendrites (MAP2), the astrocytic marker (GFAP), and the oligodendrocytic marker (O4). Scale bar, 50 µm.

NPCs were differentiated from hiPSCs as described previously[25]. and characterized via immunocytochemistry (ICC) and flow cytometry. ICC staining demonstrated the presence of neuronal lineage and stem cell markers, including PAX6, Nestin, and SOX2 (Figure 2b). Flow cytometry analysis corroborated ICC findings, revealing that over 60% of the cell population expressed all three markers (Figure 2c). Neurospheres were evaluated based on their physical integrity, viability, differentiation into neuronal and glial lineages, and functional network development. Physical properties, viability, and necrotic core formation were assessed biweekly. The spheroids maintained a compact structure throughout the eight-week culture period (Figure 2d). Spheroid diameter increased from approximately 400 µm (week 2) to 700 µm (week 8), with larger variability observed at later stages. Circularity remained consistent until week 6 but declined by week 8, suggesting structural changes at later stages (Figure 2e).

Cell viability was quantified by ATP levels, which indicated increased cell death after 6 weeks (Figure 2g); however, no necrotic core was detected (Figure 2f). Functional maturation of the neuronal network was assessed by measuring mean firing rate (MFR) and the emergence of network burst activity. MFR increased during early development but declined after 6 weeks, coinciding with the onset of network bursts in 6-week-old spheroids. These changes collectively suggest the establishment of a more mature neuronal network around week 6 (Figure 2h and Figure S2).

To evaluate cellular diversity within the spheroids, ICC protocols were optimized for dense 3D tissue, incorporating tissue-clearing techniques to enhance imaging quality. Differentiation into neuronal and glial populations was confirmed through ICC staining for axonal markers (TUBB3, NFH), dendritic markers (MAP2), astrocytic markers (GFAP), oligodendrocytic marker (O4), and cortical layer-specific markers (SATB2, CTIP2). Astrocytes emerged within 2 weeks and increased in abundance over time. By week 6, neurospheres expressed markers of both upper and lower cortical layers, indicating advanced phenotypic complexity (Figure 2i).

### NeuroMPS enables electrophysiological assessment of pharmacodynamic effects

Network activity in the brain is regulated by both inhibitory and excitatory neurons. Disruption of either neuronal subtype can lead to widespread alterations across the neural network. To benchmark the NeuroMPS model, we applied various pharmacological blockers and evaluated the resulting electrophysiological alterations with respect to alignment with expected pharmacodynamic effects.

Since network bursts increased until day 9 post-embedding and then slowly declined (Figure S2), electrophysiological modulations were investigated at day 9 post-embedding. Data were acquired after 10 minutes of compound incubation and normalized to baseline spontaneous activity (Figure 3a). GABAergic (inhibitory) neurons were targeted using picrotoxin (PTX)[26] and bicuculline (BIC)[27], both of which induced seizure-like activity in excitatory neurons. Blocked inhibitory control led to a marked increase in burst frequency rate (BFR) (Figure 3b, c). Following PTX treatment, tetrodotoxin (TTX), a sodium channel blocker, was applied to the neurospheres. Sodium channels are essential for action potential generation, and their blockade effectively silenced the signaling cascade. Accordingly, a dramatic reduction in electrical activity was recorded (Figure 3b).

**Figure 3.**
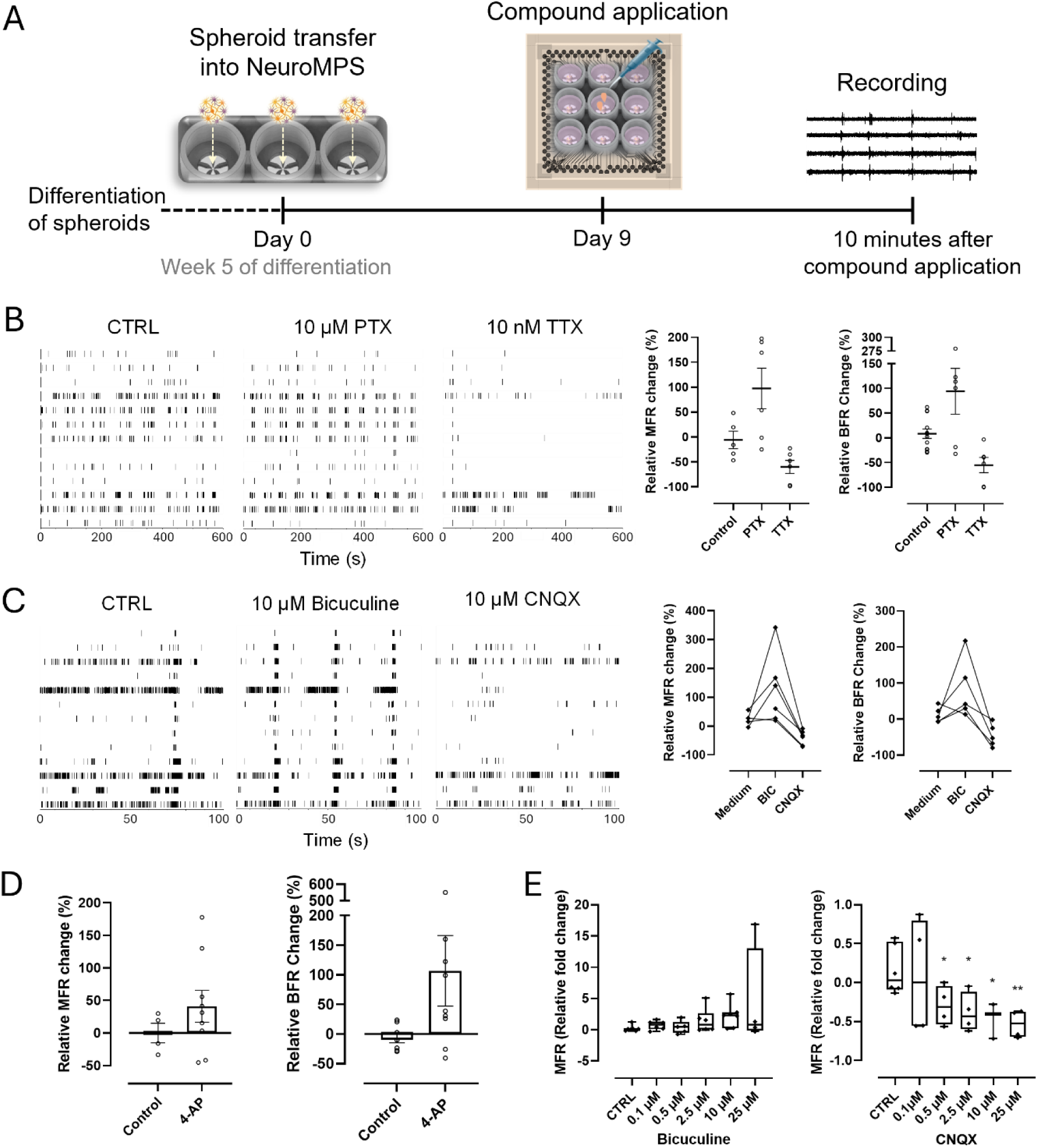
Modulation of neuronal network activity. **a** Schematic overview of compound application and electrophysiological recording workflow. **b** Representative raster plots showing neuronal activity after treatment with PTX and followed by TTX, where each line corresponds to a single spike (left). Change in MFR and BFR is shown (right); horizontal lines indicate mean values and whiskers represent individual recordings (N = 5–8). **c** Raster plots depicting network responses to BIC and CNQX (left). Corresponding relative changes in MFR and BFR are shown (right); lines connect paired measurements from the same samples (N = 5). **d** Effects of 4-AP on MFR and BFR; bar graphs represent mean values with whiskers indicating individual recordings (N = 7–9). **e** Dose–response analysis of NeuroMPS sensitivity to increasing concentrations of BIC and CNQX. Box plots indicate median and interquartile ranges; data are compared to control using Welch’s t-test, with asterisks denoting statistical significance.

The GABA antagonist BIC was further paired with cyanquixaline (CNQX) to dissect contributions from excitatory and inhibitory activity. After BIC-induced seizure-like bursting, CNQX was introduced to block all excitatory pathways except those mediated by N-methyl-D-aspartate (NMDA) receptors. While BIC inhibited the inhibitory pathways, CNQX suppressed most excitatory signaling. This dual blockade eliminated seizure-like activity but also significantly dampened overall network function, as evidenced by reductions in mean firing rate (MFR) and burst frequency rate (BFR) (Figure 3c).

To assess dose-response sensitivity, BIC and CNQX were tested at five concentrations (0.1 µM, 0.5 µM, 2.5 µM, 10 µM, and 25 µM). Changes in network activity were quantified and compared to control conditions using Welch’s t-test. BIC elicited a statistically significant response only at 10 µM, whereas CNQX produced significant effects at all concentrations except 0.1 µM (Figure 3e).

In addition to these antagonists, 4-aminopyridine (4-AP), a potassium channel blocker, was used to evaluate NeuroMPS further. Potassium channels play a critical role in repolarizing the neuronal membrane during action potential propagation. Their blockade induces neuronal stress and promotes a seizure-like phenotype by impairing repolarization. Neurospheres were treated with 30 µM 4-AP, and their responses were recorded following a 10 min incubation period. Raster plots revealed a transition to burst trains (Figure S3). Potassium channel inhibition led to increased BFR and elevated the proportion of bursts within network bursts, indicating heightened excitability across the network (Figure 3d).

### NeuroMPS enables metabolic, morphological, and functional evaluation of neurotoxicity

Finally, a proof-of-concept neurotoxicity assay was conducted using rotenone, a pesticide known to inhibit mitochondrial complex I, induce oxidative stress, and ultimately trigger apoptosis. Neurospheres differentiated for 6 weeks were exposed to rotenone, and their effects were assessed at multiple time points using a dynamic toxicity assay, morphological analyses, ATP measurements, and electrophysiological recordings(Figure 4a)

**Figure 4.**
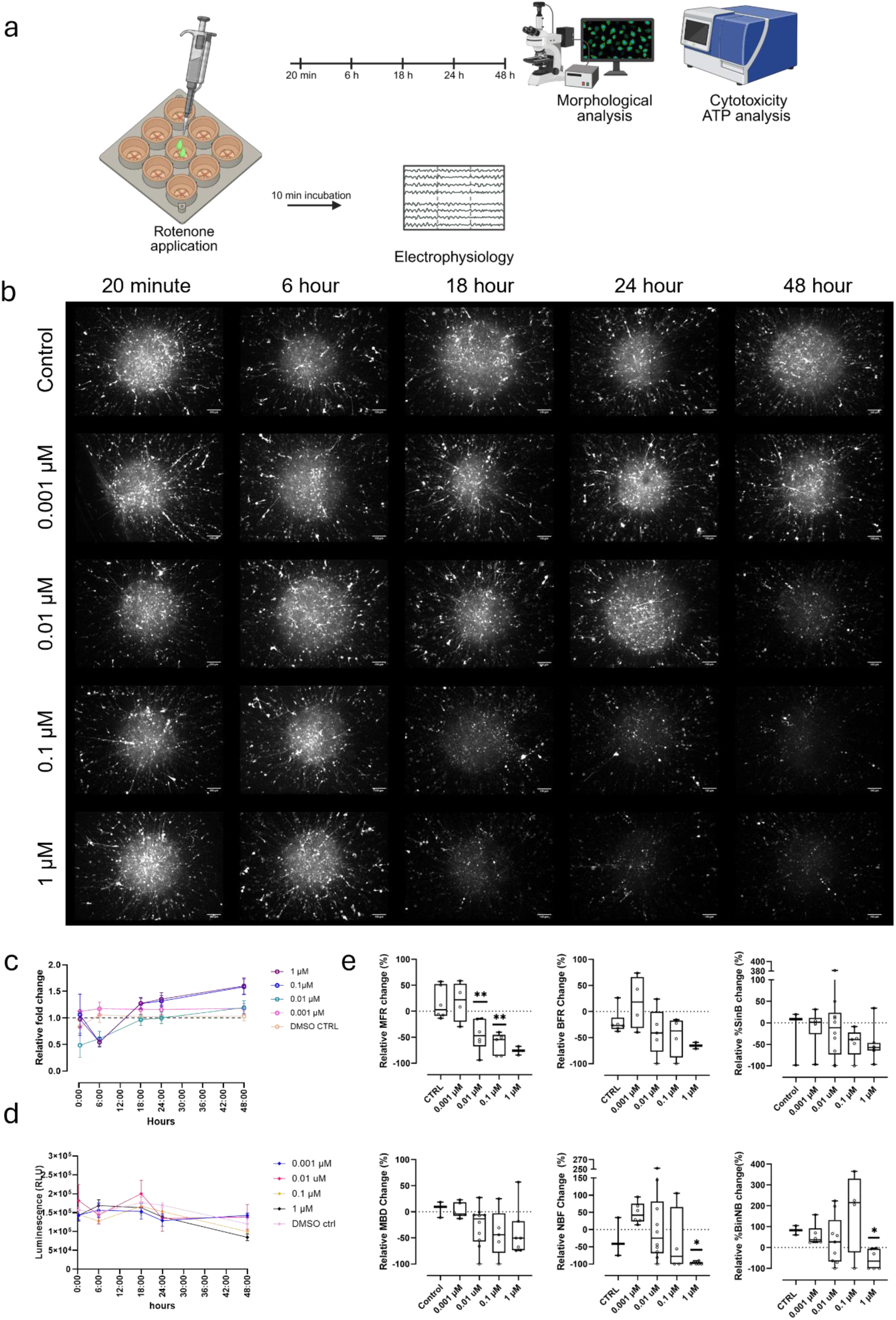
Assessment of neurosphere responses to neurotoxic exposure. **a** Schematic overview of the experimental design. Created with BioRender.com **b** Morphological assessment of KOLF2.1J neurospheres following rotenone treatment. GFP-expressing neurospheres were imaged at 20 min, 6 h, 18 h, 24 h, and 48 h post-treatment. Z-stack imaging across a 300 µm thickness was acquired and presented as maximum intensity projections. Scale bars, 100 µm. **c** Cytotoxicity was evaluated using a DNA-binding cyanine dye (CellTox^TM^ Green) to label dead cells; fluorescence intensity was quantified at 0, 6, 18, 24, and 48 h and is shown as fold change relative to time 0 (N = 6). **d** Cell viability was assessed by ATP levels at the same time points (0–48 h), demonstrating a time-dependent decrease following rotenone exposure (N = 6). **e** Electrophysiological responses of KOLF2.1J neurospheres to rotenone were recorded 10 min after compound application, with 10-min recording. Network activity parameters, including MFR, BFR, percentage of spikes in bursts (%SiNB), mean burst duration (MBD), network burst frequency (NBF), and percentage of bursts in network bursts (%BiNB), were analyzed (N = 3–9). Box plots indicate medians and interquartile ranges, with whiskers representing individual recordings and extrema. Data were compared to control using the non-parametric Kolmogorov–Smirnov test; asterisks denote statistically significant differences.

Morphological analysis was enabled by transducing neurospheres with green fluorescent protein (GFP). Imaging was performed at 20 min, 6 h, 18 h, 24 h, and 48 h following treatment. Neurite dissociation was first observed at 18 hours in wells treated with higher concentrations of rotenone (1 µM and 0.1 µM) (Figure 4a). High-magnification imaging further illustrated the progressive neurite degeneration over time (Figure S4)

To assess cytotoxicity, a microscopy-based approach using the CellTox^TM^ Green Cytotoxicity assay was employed. The earliest cytotoxic responses were detected after 18 hours at higher concentrations of rotenone (0.1 and 1 µM) (Figure 4b). Cell viability was further evaluated by quantifying ATP contents, revealing a measurable reduction in ATP only after 24 hours in all concentrations (Figure 4c).

To investigate rotenone’s impact on neuronal network activity, we used multimodal recordings via MEAs. Unlike morphological and metabolic endpoints, network disruptions occurred rapidly, with electrical circuit impairments detected within 10 minutes of treatment. The analyses revealed concentration-dependent disruptions, with higher doses producing more pronounced effects. Furthermore, rotenone markedly reduced network synchronicity at elevated concentrations, as reflected in NBF and %BinNB parameters. Even the lowest concentration caused minor, though non-significant, desynchronization (Figure 4d).

## Discussion

The NeuroMPS was engineered to meet the specific micro-physiological requirements of neurospheres, bridging a longstanding gap between 3D culture formats and advanced electrophysiological readouts. In this study, we developed a platform that incorporates three key design features: an MEA, CMEs, and custom quartz wells equipped with pins to accommodate neurospheres. This configuration ensures stable, long-term recordings while maintaining the structural integrity of the neurospheres. A robust differentiation method modified from Pamies et al.[16] provided neuronal and astroglial populations within the neurospheres. Given astrocytes’ essential role in shaping synaptic homeostasis and coordinating network function in vitro[28–30], their consistent presence likely underpins the reliability of electrophysiological outcomes. Functional maturation followed an expected trajectory: spontaneous network activity was established by six weeks, suggesting that NeuroMPS supports the coordinated emergence of circuit-level dynamics. CMEs provided a non-invasive electrophysiological readout unlike the other platforms reported before.

The fully glass-based architecture of NeuroMPS mitigates the small-molecule absorption associated with PDMS platforms.[31, 32], enabling more reliable pharmacological interrogation of neurosphere networks. Pharmacological profiling demonstrated that the system resolves dynamic shifts in excitatory–inhibitory balance: GABA receptor antagonists elicited seizure-like discharges that were subsequently suppressed by sodium channel blockade, while CNQX abolished activity, confirming functional excitatory and inhibitory connectivity. Dose-dependent CNQX responses and the pronounced hyperexcitability induced by 4-AP further illustrate the platform’s capacity to model diverse network states and synaptic perturbations. Neurotoxicity testing with rotenone revealed that electrophysiological impairments emerge within minutes, preceding metabolic and morphological changes that appear only after prolonged exposure and at higher concentrations, underscoring the sensitivity of network-level readouts as early indicators of neurotoxic stress. Together with its transparent, ANSI/SLAS-compatible design that supports parallel functional, metabolic, and morphological assays and effluent collection, NeuroMPS provides a scalable and human-relevant platform for neurotoxicity screening, pharmacological characterization, and mechanistic studies of network dysfunction.

Although the platform does not yet reach the throughput of plate-based screening systems[15, 33], it accommodates nine wells per device, enabling parallel recordings while preserving the advantages of 3D microphysiological culture. The current configuration lacks perfusion; thus, nutrient depletion and waste accumulation follow the kinetics of conventional static formats[34]. A key next step will be to render the system immunocompetent by incorporating microglia. However, while the glass-based architecture effectively addresses compound absorption, glass surfaces are known to activate immune cells, indicating that alternative materials may be required for future iterations. These considerations outline a clear trajectory for advancing NeuroMPS toward more physiologically integrated and higher-throughput applications.

## Methods

### Microfabrication

NeuroMPSs were fabricated using a protocol previously described[7], with minor modifications to enable functional neurosphere culture. Culture modules were produced from fused silica using selective laser etching. Custom MEAs were obtained from NMI TT (Reutlingen, Germany), and CMEs were patterned on gold electrodes using epoxy-based photoresists as previously described[7]. Briefly, SU-8 2002 (Kayaku, USA) was spin-coated to a thickness of approximately 3 µm (10 s at 500 rpm followed by 30 s at 1000 rpm) to form microtunnels. A 5-min soft bake at 95 °C (hotplate, temperature ramped from/back to room temperature (RT)) was performed before exposure. Structures were patterned using a SÜSS MA6 mask aligner (i-line, soft contact, 120 mJ/cm). A post-exposure bake identical to the soft bake cross-linked the exposed SU-8. Development was carried out in mr-Dev 600 (microresist technologies) for 15 s, followed by an isopropanol rinse. An intermediate hard bake at 150 °C ensured sufficient cross-linking to prevent overdevelopment during subsequent processing steps. For the second layer, ADEX TDFS A20 (Micro Resist Technology GmbH, Germany) was laminated using a GMP Photonex@325 pouch laminator (3 mm/s, 75 °C). After removing the carrier foil, the samples were exposed (600 mJ/cm, i-line, 10 µm proximity) and baked for 1 h at 65 °C (ramped from RT with 30 % power). Development was performed in cyclohexanone for 8 min, followed by isopropanol rinsing and nitrogen drying.

The final hard bake was conducted at 170 °C for 30 min (0.5 °C/min ramp from/back to RT) to complete cross-linking, ensuring chemical stability and preventing toxicity. The microfluidic chip was then coated with APTES (3-aminopropyl)-triethoxysilane) (440140, Sigma Aldrich), aligned to the MEA using a fine placer, and bonded with epoxy adhesive (EPO-TEK 301-2FL, Epoxy Technology). The adhesive was cured at 80 °C for 3 h (0.5 °C/min ramp from/back to RT).

Before use, assembled devices were cleaned in 1 % Tergazyme (Z273287, Sigma-Aldrich) solution for 2 h, followed by thorough rinsing with bi-distilled water. For reuse, NeuroMPSs were cleaned after each experiment by overnight incubation in 1 % Tergazyme solution at RT, followed by extensive washing in bi-distilled water overnight at RT, provided they remained structurally and functionally intact.

### Neurosphere formation

NPCs were derived from the KOLF2.1J iPSC line (JIPSC1000, The Jackson Laboratory, CT, USA) at Microorgano Lab (Tübingen, Germany) following a protocol adapted from Reinhardt et al. [25] For NPC expansion, culture plates were coated overnight at 4 °C with 5 µg/ml laminin (A29248, Thermo Fisher Scientific) in DPBS (+/+). NPCs were seeded at a density of 5 × 10⁴ cells/cm^2^ and maintained in NPC culture medium (Table S1) at 37 °C in a humidified atmosphere with 5 % CO_2_. The medium was refreshed every other day until cultures reached ∼80% confluence. The neurosphere generation protocol was modified from Pamies et al. [16] Briefly, to generate uniform neurospheres, AggreWell™ 800 plates (34815, STEMCELL Technologies) were pre-treated with Anti-Adherence Rinsing Solution (07010, STEMCELL Technologies) by adding 500 µl per well, followed by centrifugation at 1300 × g for 5 min. Wells were subsequently rinsed twice with NPC culture medium. NPCs were dissociated using StemPro™ Accutase™ Cell Dissociation Reagent (A1110501, Gibco) for 5 min at 37 °C with 5% CO_2_. Accutase activity was quenched with NPC culture medium and centrifuged at 300 × g for 5 min. Cells were resuspended in NPC culture medium and seeded into AggreWell™ 800 plates at a density of 1.5 × 10⁶ cells per well, followed by centrifugation at 100 × g for 3 min to promote aggregation. After homogeneous cell distribution was confirmed, plates were incubated for 48 h at 37 °C with 5 % CO_2_.

After 48 h, the medium was replaced with spheroid culture medium (Table S2), and neurospheres were transferred to 6-well plates. Each well contained 2 ml of spheroid culture medium, and plates were maintained on an orbital shaker at 90 rpm under standard culture conditions (37 °C, 5 % CO_2_). 75% of the medium was replaced every other day for 3 weeks, after which the cultures were transitioned to spheroid maturation medium (Table S3).

### Flow cytometry

The quality of NPCs was evaluated using flow cytometry. Cells were initially stained with extracellular antibodies/dyes (Table S4) in DPBS (−/−) for 10 min at RT in the dark. The FOX-P3 Buffer Kit (130-093-142, Miltenyi Biotec) was used for intracellular staining (Table S4). Cells were fixed and permeabilized with the kit’s permeabilization/fixation solution for 30 minutes at 4 °C in the dark, then incubated with the intracellular antibody solution in the kit’s permeabilization solution for another 30 minutes at 4 °C. After each step, cells were centrifuged at 300 × g for 4 minutes and washed with AutoMACS running buffer (130-091-221, Miltenyi Biotec). The cells were then resuspended in AutoMACS running buffer and maintained at 4 °C until acquisition. An unstained control was included in each experiment. Samples were acquired using the BD LSRFortessa™ Cell Analyzer (BD Biosciences), and data were analyzed using FlowJo software (version 10, Waters^TM^ Biosciences).

### Culturing in NeuroMPS

5 week-old neurospheres were embedded in Matrigel (356230, Corning) diluted to 75 % (v/v) in spheroid maturation medium. Individual neurospheres were transferred into the NeuroMPS and positioned on top of pins within the Matrigel matrix, one neurosphere per well. The NeuroMPSs were incubated at 37 °C for 10 min to allow gel polymerization. Following gelation, spheroid maturation medium was added to each well, and cultures were maintained under standard conditions (37 °C, 5 % CO_2_).

### Immunocytochemistry

The neurospheres were fixed in a solution containing 4 % PFA (P6148, Sigma-Aldrich) and 4% sucrose (S0389, Sigma-Aldrich) in PBS for 1 h at RT, then washed 3 times for 5 minutes each in DPBS (14190144, Thermo Fisher Scientific, MA, USA). The spheroids were dehydrated using a series of solutions (Table S5). Spheroids were sequentially incubated in Solution 1 for 15 minutes, followed by Solution 2 for 15 minutes, and then Solution 3 for an additional 15 minutes. The spheroids were subsequently returned to Solution 1 for another 15 minutes, then washed in DPBS for 15 minutes. After the washing step, spheroids were permeabilized with Solution 4 for 30 minutes. Then, the spheroids were transferred into the penetration buffer (HSB-BK, Visikol, NJ, USA) and incubated for 30 minutes at 300 rpm. All steps were performed at RT up to this point. The spheroids were then incubated in blocking buffer (HSB-BK, Visikol, NJ, USA) at 37 °C for 1 h at 300 rpm. After blocking, the spheroids were incubated with primary antibodies (Table S6) prepared in the antibody buffer (HSB-BK, Visikol, NJ, USA) overnight at 37 °C, at 300 rpm, followed by an additional 48 h at 4 °C (no mixing). They were then washed 3 times for 15 minutes in the washing buffer (HSB-BK, Visikol, NJ, USA) at 37 °C with mixing at 300 rpm. Secondary antibodies and the nuclear dye (Table S7) were prepared in the antibody buffer, and neurospheres were immersed in this solution. The spheroids were then incubated at 37 °C for 90 minutes at 300 rpm. The spheroids were washed three times for 15 minutes in washing buffer at 37 °C and 300 rpm to remove unbound secondary antibodies. Then, the spheroids were brought to RT, the washing buffer was replaced with DPBS, and spheroids were incubated at RT for 10 minutes, and then cleared in Scale S4 solution[35] (Table S8) at 4 °C for 48-72 hours. Before imaging, the neurospheres were mounted in SlowFade^TM^ Diamond Antifade mountant (S36967, Thermo Fisher Scientific, MA, USA). The images were acquired using spinning-disk confocal microscopy (Cell Observer^®^ with spinning disk head, Zeiss).

### AAV1-EGFP Transduction

One day after plating in NeuroMPS, 5-week-old neurospheres were transduced with adeno-associated virus serotype 1 (AAV1) carrying a transgene encoding enhanced green fluorescent protein (eGFP) under the control of the human Synapsin-1 (hSyn) promoter (pAAV-hSyn-EGFP, #50465-AAV1, Addgene). A multiplicity of infection (MOI) of 10⁵ was used, and eGFP expression was monitored daily by fluorescence microscopy. Pharmacological compounds were applied 7 days after transduction.

### Physical characterization of the neurospheres

Neurospheres were assessed for circularity and diameter over time. Bright-field images were acquired every two weeks using an EVOS FL digital fluorescence microscope (ThermoFisher Scientific). For each well of a 6-well plate, five randomly selected regions were imaged. Image analysis was performed using a custom ImageJ macro. Images were first converted to binary and subsequently transformed into masks. The “Analyze Particles” function was applied using the following parameters: size = 30,000–∞ µm², include holes, and exclude on edges. Neurosphere shape was approximated as circular, and the diameter was calculated from the measured area (A) using the formula:

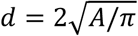

The circularity of the neurospheres was calculated using the formula below, where A indicated area and P indicated perimeter:

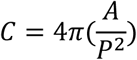

### Live/Dead Staining

Neurospheres were transferred into 96-well plates for the assay. They were stained with 1:2000 CellTox™ Green (G8741, Promega GmbH) and 20 µM Hoechst (62249, Thermo Fisher Scientific) in media. The dye solution was prepared in neurosphere culture medium, and the spheroids were incubated in the staining solution for 30 minutes at 37 °C with 5 % CO_2_.

Following incubation, neurospheres were imaged using spinning-disk confocal microscopy (Cell Observer® with spinning disk head, Zeiss).

### Cell Viability Assay

Neurospheres were transferred into a 96-well plates prior to the assay. The CellTiter®-Glo 3D Cell Viability Assay (G7570, Promega GmbH) was performed according to the manufacturer’s instructions. Briefly, the reagent was added to each well at a 1:1 ratio with the existing amount of media. Plates were then shaken at 800 rpm for 5 min in a thermomixer (ThermoMixer C, Eppendorf) to enhance cell lysis, followed by a 25-min incubation at RT in the dark. Luminescence was subsequently measured using a TECAN Spark plate reader according to the protocol provided by Promega.

### Dynamic Cytotoxicity Assay

5 week-old neurospheres were transferred into a 96-well plates, stained with 1:2000 CellTox™ Green in media for 30 minutes, and then treated with rotenone at the concentrations of 0.001, 0.01 µM, 0.1 µM, and 1 µM The neurospheres were imaged using the ImageXpress Micro-Confocal High-Content Imaging System (Molecular Devices) at 6 h, 18 h, 24 h, and 48 h. For analysis, intensity values were normalized to untreated controls.

### Electrophysiological recordings

The electrical activity of neurospheres was recorded using an MEA2100-System (Multi Channel Systems MCS GmbH, Germany) at defined time points, enabling simultaneous acquisition from 120 channels at a maximum sampling rate of 50 kHz per channel. The system contained an integrated filter amplifier with adjustable gain and bandwidth controlled through the acquisition software (MC Rack). Spontaneous neuronal network activity of neurospheres was recorded nine days after plated in the NeuroMPS. All experiments were performed under continuous monitoring of temperature, CO_2_, and relative humidity using a customized MEA incubation chamber (Okolab Srl, Italy). Recordings began 10 minutes after the chamber was closed to allow environmental conditions to stabilize. The temperature was maintained at 36 °C, relative humidity at 85%, and CO_2_ at 5%.

Spike detection, burst analysis, and network burst quantification were performed using NeuroExplorer (version 5.300). Raw signals were first filtered using a fourth-order bandpass filter (60–6000 Hz). Action potentials were detected using a threshold of T = 4.75 σ_N_, where σ_N_ represents the estimated standard deviation of background noise[36]. The minimum inter-spike interval was set to 1 ms. Burst detection was performed using the Max Interval method, as previously characterized. Network bursts within each well of the NeuroMPS were identified using a previously described custom Python script [7]. Briefly, the script calculated the median spike time for each burst on every electrode and projected these medians as single events onto a unified timeline. Network bursts were defined based on a Poisson distribution of this sequence of medians, using a surprise value of 3 and a minimum duration of 5 ms. Network burst duration was calculated as the average time between the first and last spikes of all bursts contributing to the network event. A minimum of 33.3 % of electrodes (i.e., six electrodes per well) was required for an event to be classified as a network burst.

### Compound application

Responses to PTX (1128, Tocris Bioscience), TTX (1078, Tocris Bioscience), bicuculline methchloride (BIC) (B7686, Sigma Aldrich), CNQX disodium salt hydrate (C239, Sigma Aldrich), 4-aminopyridine (4-AP) (A78403, Sigma Aldrich), and rotenone (R8875, Sigma Aldrich) were assessed after adding each compound to individual wells of the device placed inside the MEA recording system. Five percent of the culture medium was replaced with fresh medium containing test compounds at 20X the final concentration, and the mixture was carefully mixed using an electronic multichannel pipette (VIAFLO 12-channel, 0.5 – 12.5 µl, INTEGRA Biosciences). All experiments included a negative control to assess the effect of the application procedure. Recordings began 10 min after compound application and continued for 10 min.

### Statistical Analysis

Statistical analyses were performed using GraphPad Prism (version 9.3.1). Sample sizes were mentioned in the corresponding figure captions. Baseline values for each condition were first assessed for outliers using the ROUT method (Q = 5 %). Data normality was evaluated with the Shapiro–Wilk test. Depending on the distribution and experimental design, comparisons between two groups were conducted using a Student’s *t*-test, and comparisons among multiple groups were performed using one-way ANOVA. The specific statistical test used was stated in each figure caption. Error bars represented the standard error of the mean (SEM). Statistical significance was denoted as *p* < 0.05; \**p* < 0.01; \*\**p* < 0.001; ns, not significant unless stated otherwise.

## Acknowledgements

We thank Madalena Cipriano for contributions to neurotoxicology experiment design and Claudia Teufel for flow cytometry data analysis. We also thank Michael Mierzejewski for his support with microfabrication processes. This work was supported by EUROSTARS under grant agreement E!115217 (NEUROCHIP).

