## Supporting Information for "Electrophysiological profiling of hiPSC-derived neurospheres using a novel NeuroMPS with integrated electrodes"

**Table S1. NPC Culturing Medium Composition**

| | WORKING CONCENTRATION | PRODUCT INFO |
|---|---|---|
| **NEUROBASAL™ MEDIUM** | 1:1 with DMEM/F12 medium | 21103049, Gibco, Thermo Fisher Scientific, MA, USA |
| **DMEM/F-12, GLUTAMAX™ SUPPLEMENT** | 1:1 with Neurobasal™ medium | 31331028, Gibco, Thermo Fisher Scientific, MA, USA |
| **B-27™ SUPPLEMENT (50X), MINUS VITAMIN A** | 1X | 12587010, Gibco, Thermo Fisher Scientific, MA, USA |
| **N-2 SUPPLEMENT (100X)** | 1X | 17502048, Gibco, Thermo Fisher Scientific, MA, USA |
| **PENICILLIN-STREPTOMYCIN (10,000 U/ML)** | 100 U/ml | 15140122, Gibco, Thermo Fisher Scientific, MA, USA |
| **2-MERCAPTOETHANOL (50 MM)** | 50 µM | 31350010, Gibco, Thermo Fisher Scientific, MA, USA |
| **HUMAN/MOUSE/RAT BDNF, ANIMAL-FREE RECOMBINANT PROTEIN, PEPROTECH®** | 20 ng/ml | AF-450-02, Gibco, Thermo Fisher Scientific, MA, USA |
| **HUMAN FGF-BASIC (FGF-2/BFGF) (154 AA), ANIMAL-FREE RECOMBINANT PROTEIN, PEPROTECH®** | 10 ng/ml | AF-100-18B, Gibco, Thermo Fisher Scientific, MA, USA |
| **RECOMBINANT HUMAN EGF PROTEIN, CF** | 10 ng/ml | 236-EG, R&D Systems Inc., MN, USA |

**Table S2. Spheroid Culturing Medium Composition**

| | WORKING CONCENTRATION | PRODUCT INFO |
|---|---|---|
| **NEUROBASAL™ MEDIUM** | | 21103049, Gibco, Thermo Fisher Scientific, MA, USA |
| **B-27™ SUPPLEMENT (50X), SERUM-FREE** | 1X | 17504044, Gibco, Thermo Fisher Scientific, MA, USA |
| **GLUTAMAX™ SUPPLEMENT (100X)** | 1X | 35050038, Gibco, Thermo Fisher Scientific, MA, USA |
| **PENICILLIN-STREPTOMYCIN-GLUTAMINE (100X)** | 1X | 10378016, Gibco, Thermo Fisher Scientific, MA, USA |
| **HUMAN/MOUSE/RAT BDNF, ANIMAL-FREE RECOMBINANT PROTEIN, PEPROTECH®** | 10 ng/ml | AF-450-02, Gibco, Thermo Fisher Scientific, MA, USA |
| **RECOMBINANT HUMAN GDNF PROTEIN** | 10 ng/ml | 212-GD, R&D Systems Inc., MN, USA |

**Table S3. Spheroid Maturation Medium Composition**

| | WORKING CONCENTRATION | PRODUCT INFO |
|---|---|---|
| **BRAINPHYS™ MEDIUM** | | 5790, STEMCELL Technologies Inc., Germany |
| **B-27™ SUPPLEMENT (50X), SERUM-FREE** | 1X | 17504044, Gibco, Thermo Fisher Scientific, MA, USA |
| **GLUTAMAX™ SUPPLEMENT (100X)** | 1X | 35050038, Gibco, Thermo Fisher Scientific, MA, USA |
| **PENICILLIN-STREPTOMYCIN-GLUTAMINE (100X)** | 1X | 10378016, Gibco, Thermo Fisher Scientific, MA, USA |
| **HUMAN/MOUSE/RAT BDNF, ANIMAL-FREE RECOMBINANT PROTEIN, PEPROTECH®** | 10 ng/ml | AF-450-02, Gibco, Thermo Fisher Scientific, MA, USA |
| **RECOMBINANT HUMAN GDNF PROTEIN** | 10 ng/ml | 212-GD, R&D Systems Inc., MN, USA |

**Table 4. Flow Cytometry antibodies and dyes**

| | CLONE | CONJUGATE | APPLICATION | PRODUCT INFO |
|---|---|---|---|---|
| **PAX-6 ANTIBODY, ANTI-HUMAN, REAFINITY™** | REA507 | APC | ICFC | 130-123-267, Miltenyi Biotec GmbH, Germany |
| **NESTIN ANTIBODY, ANTI-MOUSE/RAT, REAFINITY™** | REA575 | PE | ICFC | 130-119-799, Miltenyi Biotec GmbH, Germany |
| **SOX2 ANTIBODY, ANTI-HUMAN/MOUSE, REAFINITY™** | REA320 | Vio Bright V423 | ICFC | 130-131-077, Miltenyi Biotec GmbH, Germany |
| **ZOMBIE AQUA™ FIXABLE VIABILITY KIT** | | | Surface | 423101, , BioLegend CNS Inc, CA, USA |

**Table S5. Dehydration Solutions**

| | SOLUTION RECIPE | PRODUCT INFO |
|---|---|---|
| **INGREDIENTS** | | |
| METHANOL | | 5.89596, Sigma-Aldrich, Merck KGaA, Darmstadt, Germany |
| DIMETHYL SULFOXIDE (DMSO) | | D2438, Sigma-Aldrich, Merck KGaA, Darmstadt, Germany |
| TRITON X-100 | | X100, Sigma-Aldrich, Merck KGaA, Darmstadt, Germany |
| **SOLUTIONS** | | |
| SOLUTION 1 | 50 % Methanol in water | |
| SOLUTION 2 | 100 % Methanol | |
| SOLUTION 3 | 20% DMSO in methanol | |

| | |
|---|---|
| SOLUTION 4 | 1 % Triton X-100 in PBS |

**Table S61. Primary Antibodies and dyes**

| | HOST | PRODUCT INFO |
|---|---|---|
| **PURIFIED ANTI-TUBULINB3 (TUBB3)** | Mouse | 801201, BioLegend CNS Inc, CA, USA |
| **ANTI-MICROTUBULE-ASSOCIATED PROTEIN 2 (MAP2)** | Chicken | NB300-213, Novus Biologicals, CO, USA |
| **ANTI-GLIAL FIBRILLARY ACIDIC PROTEIN (GFAP)** | Rabbit | GA52461-2, DAKO Omnis, Agilent CA, USA |
| **ANTI-NEUROFILAMENT HEAVY POLYPEPTIDE ANTIBODY (NFH)** | Chicken | ab4680, abcam Inc., UK |
| **ANTI-NESTIN ANTIBODY** | Mouse | MAB1259, R&D Systems Inc., MN, USA |
| **ANTI-PAX-6 ANTIBODY** | Rabbit | 901301, BioLegend CNS Inc, CA, USA |
| **ANTI-SOX2 ANTIBODY** | Rabbit | AB5603, Sigma-Aldrich, Merck KGaA, Darmstadt, Germany |

**Table S7. Secondary Antibodies**

| | REACTIVITY | HOST | PRODUCT INFO |
|---|---|---|---|
| **ALEXA FLUOR 488- CONJUGATED AFFINIPURE DONKEY ANTI-MOUSE IGG** | Mouse | Donkey | 715-545-150, Dianova GmbH, Germany |
| **GOAT ANTI-RABBIT IGG (HEAVY CHAIN), SUPERCLONAL™ RECOMBINANT SECONDARY ANTIBODY, ALEXA FLUOR™ 555** | Rabbit | Goat | A27039, Invitrogen, Thermo Fisher Scientific, MA, USA |
| **ALEXA FLUOR 647- CONJUGATED AFFINIPURE DONKEY ANTI-CHICKEN IGG** | Chicken | Donkey | 703-605-155, Dianova GmbH, Germany |
| **4′,6-DIAMIDINO-2-PHENYLINDOLE DIHYDROCHLORIDE (DAPI)** | | | D8417, Sigma-Aldrich, Merck KGaA, Darmstadt, Germany |

**Table S8. The Composition of Scale S4 Solution**

| | CONCENTRAT ION | PRODUCT INFO |
|---|---|---|
| **D-SORBITOL** | 40 % (w/v) | S3889, Sigma-Aldrich, Merck KGaA, Darmstadt, Germany |
| **GLYCEROL** | 10 % (w/v) | G2025, Sigma-Aldrich, Merck KGaA, Darmstadt, Germany |
| **UREA** | 4 M | 51456, Sigma-Aldrich, Merck KGaA, Darmstadt, Germany |
| **TRITON X-100** | 0.2 % (w/v) | X100, Sigma-Aldrich, Merck KGaA, Darmstadt, Germany |
| **DMSO** | 15 % (w/v) | D2438, Sigma-Aldrich, Merck KGaA, Darmstadt, Germany |
| **ULTRAPURE WATER** | | |

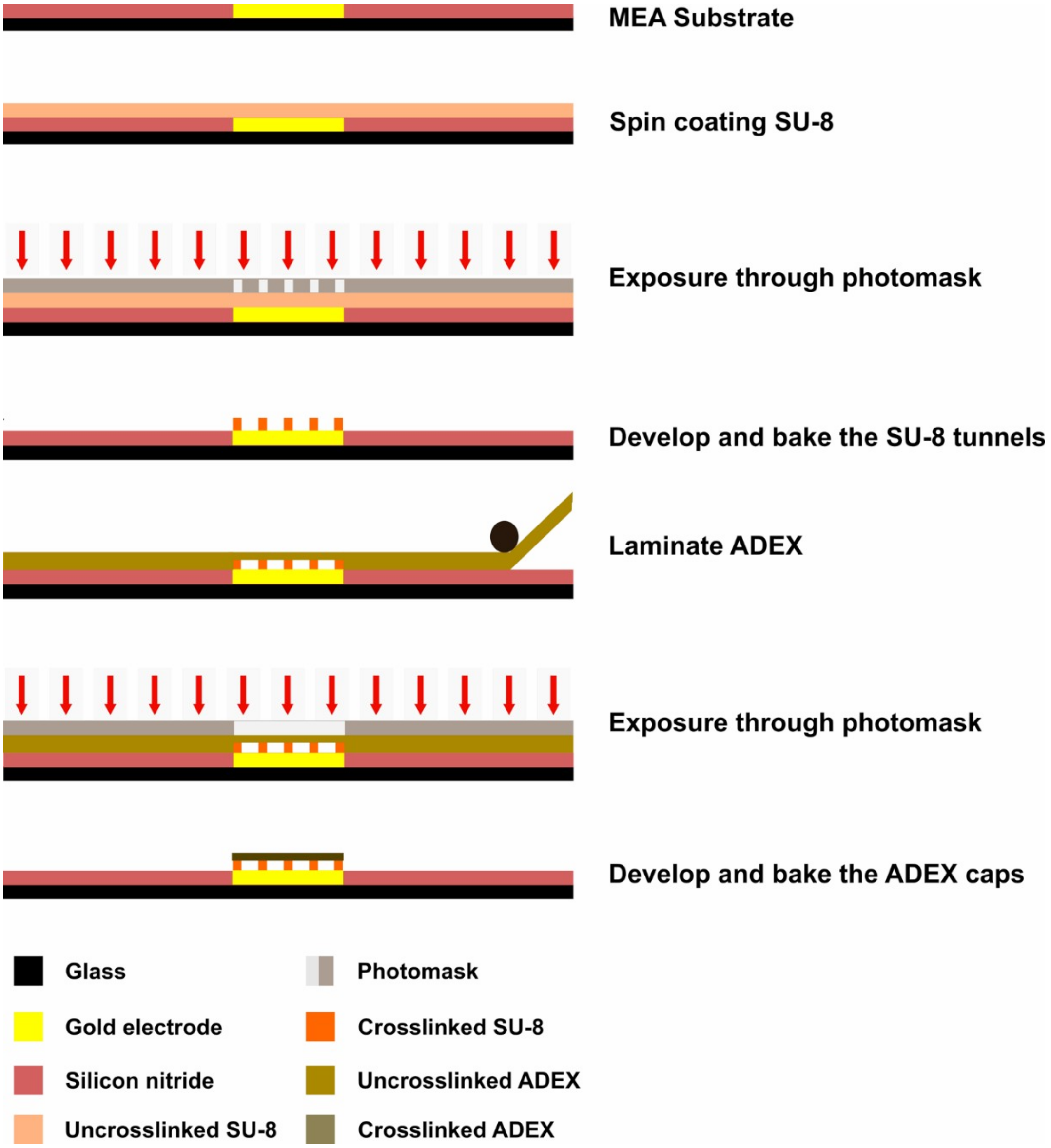


**Figure S1** Photolithography process of CME fabrication

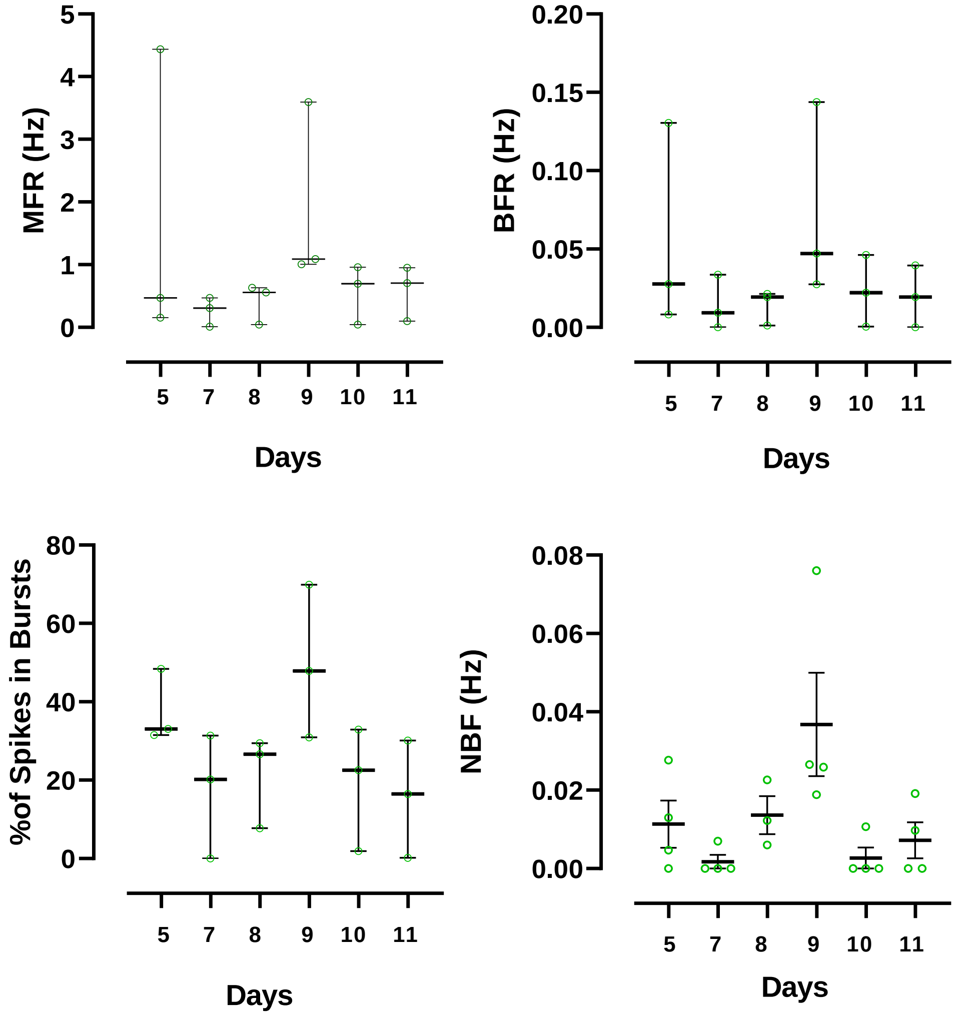


**Figure S2** Activity development of neurospheres in NeuroMPS. The neurospheres are transferred into neuroMPS at week 5, and the recordings are performed on days 5, 7, 8, 9, 10, and 11. The data is analyzed for mean firing rate (MFR), Burst frequency rate (BFR), % of spikes in Bursts (%SinB), and network burts frequency (NBF) . N= 3-5.

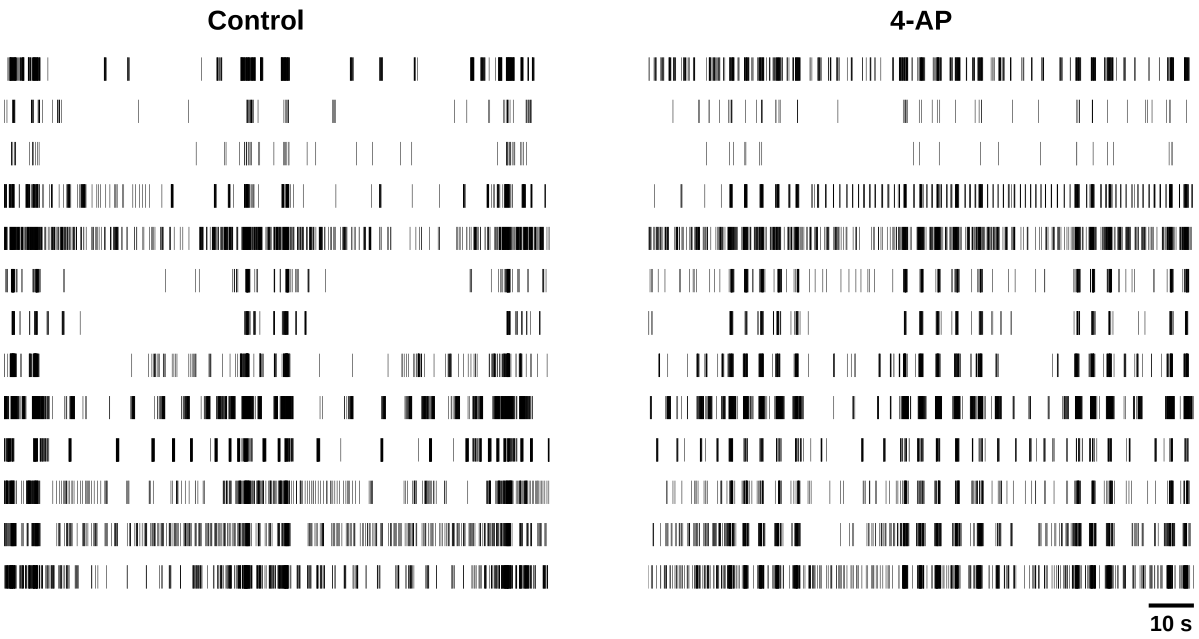


**Figure S3** Raster plots for response to 4-AP. Each vertical line represents one spike. Each horizontal train of lines represents the recordings from individual electrodes.

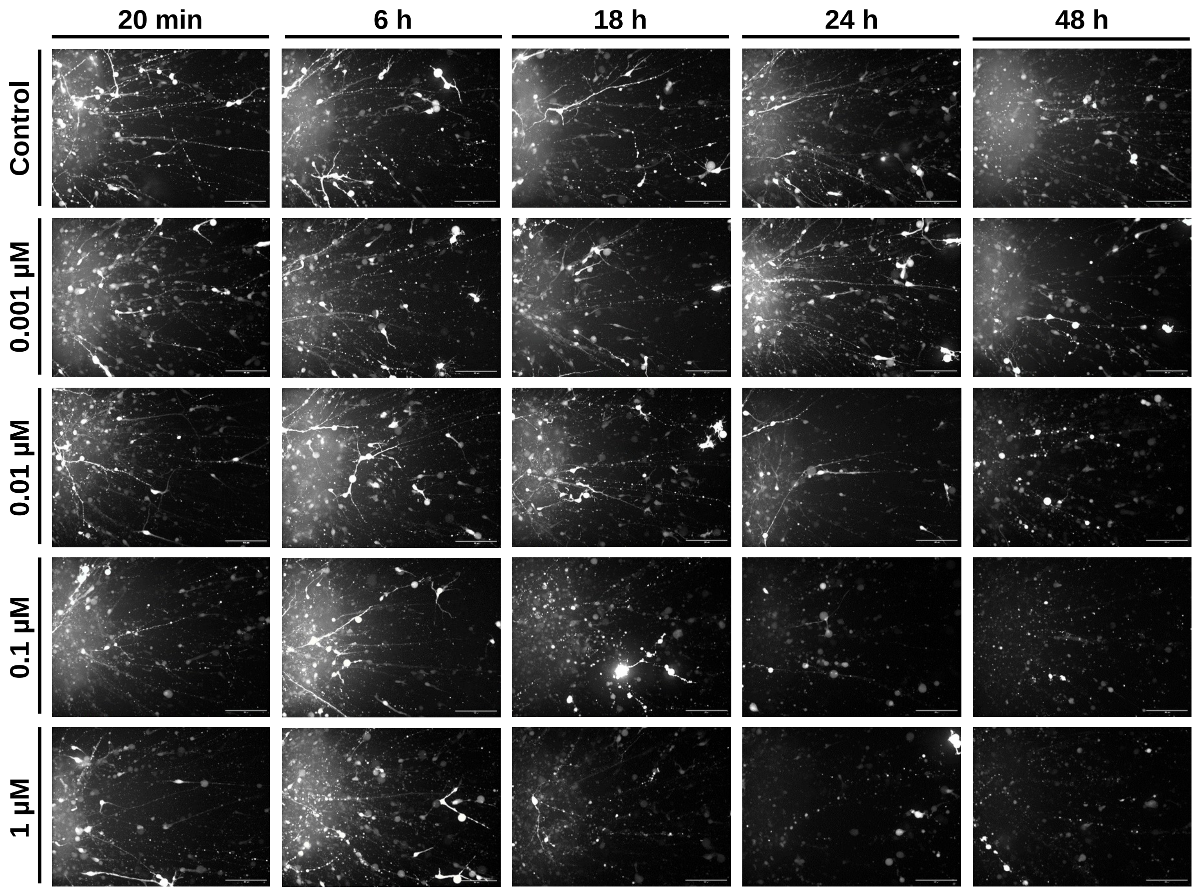


**Figure S4** Neurite focused morphological assessment of KOLF2.1J neurospheres after rotenone treatment. GFP expressing neurospheres are treated with rotenone and imaged after 20 minutes, 6 hours, 18 hours, 24 hours and 48 hours. Z-stack imaging is employed through 300 µm thickness and images are demonstrated as MIP. Scale bars, 100 µm.
